# Research on crop phenotype prediction using SNP context and whole-genome feature embedding

**DOI:** 10.1101/2025.04.08.647762

**Authors:** Huan Li, Yunpeng Cui, Tan Sun, Ting Wang, Zhen Chen, Chao Wang, Wenbo Bian, Juan Liu, Mo Wang, Li Chen, Jinming Wu, Jie Huang

**Affiliations:** Institute of Agricultural Information, Chinese Academy of Agricultural Sciences (CAAS), Beijing 100081, China; Key Laboratory of Agricultural Big Data, Ministry of Agriculture and Rural Affairs, Beijing, China; Collaborative Innovation Center of Henan Grain Crops, Henan Agricultural University, Zhengzhou 450046, China; Zibo Digital Agriculture and Rural Research Institute, Zibo 255000, China

**Keywords:** genomic prediction, DNABert-2, partial least squares, principal component analysis, SNP-context feature embedding, whole-genome feature embedding

## Abstract

Modern agriculture demands precise genomic prediction to accelerate elite crop breeding, yet traditional genomic prediction approaches, such as genomic best linear unbiased prediction (GBLUP) and Bayesian methods, focus primarily on the cumulative effect of individual SNPs, thus neglecting the concerted influence that the surrounding sequence context has on the phenotype. To overcome these limitations, we propose two novel feature embedding modes (SNP-context and whole-genome) based on DNABERT-2, a cross-species genomic foundation model that uses self-attention mechanisms and transfer learning to automatically identify conserved sequence features across diverse evolutionary lineages without prior biological assumptions. The whole-genome feature embedding aggregates genomic information at a global scale by pooling vectors from chunked sequences processed by DNABERT-2, whereas the context feature embedding captures local information by directly encoding variable-length (500--3000 bp) sequences centered on target SNPs. To reduce noise in the high-dimensional feature embeddings, we employed principal component analysis (PCA) and partial least squares (PLS) to project the features into a lower-dimensional space. We generated two kinds of feature embedding for three crop datasets (rice413, rice395, and maize301), investigated the impact of 500--3000 bp flanking SNP contexts on phenotypic prediction, and compared prediction accuracy variations across algorithms at 4--768 feature dimensions among the PCA, PLS, and no dimensionality reduction strategies. The results demonstrate that machine learning (ML) algorithms operating under the SNP-context embedding mode achieve greater accuracy and lower mean absolute errors (MAEs) than traditional SNP features do at specific context lengths, particularly for traits with low-to-moderate heritability (h^2^∈(0.2, 0.7]). In contrast, using whole-genome embeddings as input for ML can further improve the prediction accuracy for highly heritable traits (h^2^∈(0.7, 1.0]), even outperforming state-of-the-art deep learning models (such as DNNGP and ResGS) that rely on SNP markers. Our code is available on https://github.com/oliveSpring/Crop_DNA_Embedding.git

## Introduction

In recent years, the growth of the global population and significant climate changes have imposed substantial pressure on food production systems worldwide. On the one hand, the increasing population demands greater food supplies to combat persistent hunger; on the other hand, challenges such as global warming, water and soil resource scarcity, and frequent pest and disease outbreaks have severely disrupted crop growth and metabolism, ultimately reducing yield and quality [1–3]. According to the Food and Agriculture Organization (FAO), global food production must increase by more than 70% by 2050 to meet the needs of a population exceeding 9 billion [4, 5]. Accelerating the selection of crop varieties with high yield potential is pivotal for addressing this challenge. However, traditional breeding methods are limited by cumbersome experimental cycles, and several years or even decades are often needed to develop crop varieties with desired agronomic traits [6, 7]. The advent of high-throughput sequencing technology has significantly reduced genotyping costs and generated extensive high-density genetic markers, thereby accelerating the shift in crop breeding methodologies from empirical tradition to data-driven paradigms [8–10]. Genomic selection (GS), a method that employs whole-genome markers to predict genomic estimated breeding values (GEBVs), has revolutionized animal and crop breeding by enabling early-stage selection [11–16]. Nevertheless, the genetic gain achievable through GS is intrinsically limited by the accuracy of genomic prediction (GP) models. Improving GP accuracy can effectively shorten breeding cycles and reduce associated costs [17–19]. A fundamental challenge in GP arises from the high dimensionality of genomic data, where the number of single-nucleotide polymorphism (SNP) (P) markers vastly exceeds the available sample size (N), which is the so-called “big P, small N” problem. This high dimensionality (P >> N) not only substantially increases the risk of overfitting but also poses significant challenges for robustly estimating marker effects and building generalizable predictive models [17, 20]. Furthermore, linkage disequilibrium (LD) between SNPs introduces multicollinearity, which in turn increases the complexity of modeling [21]. To alleviate these issues, a subset of relevant SNP markers was selected for use in place of genome-wide markers [22]. Although whole-genome association studies (GWASs) effectively identify genetic loci associated with complex traits, pinpointing the most predictive markers for specific phenotypes remains challenging owing to the polygenic architecture of agronomic traits, confounding factors such as population stratification, and limited generalizability of marker‒trait correlations across genetically distinct cohorts.

Current genomic prediction methodologies in plant sciences focus mainly on statistical modeling and ML approaches. Statistical models include classical best linear unbiased prediction (BLUP) and its variants, such as genomic BLUP (GBLUP) [23], ridge regression BLUP (RR-BLUP) [24, 25], and single-step GBLUP [26]. A key limitation of these BLUP-based models is their assumption of a uniform genetic architecture, where all the markers are treated as having equal effect variance. This renders them less effective for traits controlled by a few major genes. While Bayesian methods, such as BayesA, BayesB, BayesCπ, and Bayes Lasso [27–29], offer more flexibility by using priors that allow for marker-specific variances, their main shortcomings lie in being computationally intensive and, more critically, remaining within a linear framework—thus limiting their ability to capture nonadditive effects such as epistasis. In contrast, ML algorithms such as support vector regression (SVR), random forest (RF), and gradient boosting regression (GBR) [30–32] serve as powerful alternatives. The primary strength lies in their inherent ability to capture complex, nonlinear relationships without relying on strict distributional assumptions. As a branch of ML, deep learning takes this capability a step further by automatically learning hierarchical features directly from genome-wide markers [33]. A representative example is DeepGS, which employs a convolutional neural network (CNN) with an 8-32-1 architecture to predict phenotypes from genotypes [34]. DNNGP is another deep neural network with three wide convolutional layers that can predict quantitative traits from multi-omics data and surpasses most genomic selection (GS) methods on four datasets [35]. Unlike DeepGS and DNNGP, SoyDNGP uses a deeper and narrower convolutional neural network (CNN) architecture with stacked small kernels for soybean genomic prediction [36]. Recently, the incorporation of residual structures and strided convolution led to the development of the 35-layer ResGS model, which achieved reliable predictions on four public datasets [32].

To date, most methods rely on genome-wide SNP markers. While these markers can strongly associate with traits, they are often used as abstract features, overlooking crucial biological contexts such as the underlying DNA sequence and local haplotype structure [30]. Moreover, SNPs represent only a fraction of genomic variations, failing to account for other types of genetic variations, such as insertions, deletions, and structural variations (e.g., copy number variations, inversions, translocations), which also significantly influence phenotypes [37]. Additionally, inaccuracies in SNP genotyping, caused by factors such as poor sample quality, insufficient sequencing depth, and the choice of alignment and variant calling algorithms, can compromise the performance of phenotype prediction models [38]. To solve these problems, we hypothesize that integrating whole-genome features or sequences adjacent to SNPs could provide more comprehensive genetic variation information, potentially enhancing the interpretability of genomic mechanisms and improving phenotype prediction accuracy.

DNA and RNA sequences are composed of four nucleotide bases (A, G, C, T, or U), analogous to the 26 letters that form words in the English language. The specific arrangement of these nucleotides encodes genetic information, much like how the sequential ordering of letters forms meaningful words and sentences. Within these sequences, distinct patterns, such as promoters, terminators, and coding regions, play crucial roles in regulating gene expression. This regulatory mechanism parallels the way specific grammatical rules govern the construction of semantically correct sentences in natural language. Furthermore, the three-dimensional folding of biological sequences enables interactions between distant sequence elements, mirroring the long-range syntactic dependencies observed between words in different positions within a sentence [39]. This similarity provides a certain scientific rationale for applying natural language processing (NLP) methods to the analysis of biological sequences.

The revolution in NLP sparked by models such as BERT [40] has inspired bioinformatics, driving the development of foundational models such as DNABERT—pretrained on the human reference genome GRCh38.p13—which have been fine-tuned for tasks such as promoter prediction, transcription factor-binding site (TFBS) identification, and splice site prediction [41]. DNABERT-2, an enhanced cross-species genomic model, improves long-sequence processing efficiency and achieves state-of-the-art performance on the GUE dataset [42]. Similarly, HyenaDNA, pretrained on a single human reference genome, leverages implicit convolutions and instruction tuning to excel in tasks such as chromatin spectrum prediction and species classification [43]. However, training such foundational models from scratch on new crop genomes is computationally demanding. Therefore, to circumvent this substantial overhead, we adopt a transfer learning strategy by employing the pretrained, cross-species DNABERT-2 to generate feature embeddings from our target crop sequences.

On the basis of the granularity of the genomic sequences to be encoded, we propose two distinct feature representation modes: (1) a whole-genome encoding mode to capture global biological patterns and (2) an SNP-context encoding mode to extract local features from the sequences flanking SNP markers. We generated feature embeddings for three crop datasets (rice413, rice395, and maize301) under two modes and employed principal component analysis (PCA) and partial least squares (PLS) for dimensionality reduction. Then, we compared the phenotypic prediction accuracy (measured by the Pearson correlation coefficient (PCC)) across different dimensions (4--768), dimensionality reduction methods (PLS, PCA, and no dimensionality reduction), and SNP context lengths. Our results demonstrate that most machine learning algorithms utilizing SNP-context embedding (at specific context lengths) outperform those based on individual SNP markers, particularly for traits with low-to-moderate heritability (h²∈ (0.2, 0.7]). Notably, for the AC trait in rice395 and the EarHT trait in maize301, the accuracy of this approach even surpassed that of models using whole-genome feature embeddings. When traits with high heritability (h²> 0.7) are predicted, conventional machine learning algorithms using whole-genome embeddings achieve significantly higher prediction accuracy and lower mean absolute error (MAE) than state-of-the-art deep learning models (e.g., DNNGP and ResGS), which rely on SNP markers.

## Materials and methods

### Plant materials

We collected publicly available datasets of genome-wide SNP markers and phenotypic traits for three crop datasets—rice413, rice395, and maize301. The rice413 and rice395 datasets were both sourced from the literature [44]. The rice413 dataset comprises 413 inbred lines of Asian rice from 82 countries, containing 44,100 SNP markers and 34 traits. For our analysis, we selected six traits with available phenotypic data: alkali spreading value (ASV; n = 403), amylose content (AC; n = 401), panicle number per plant (PNPP; n = 372), protein content (PC; n = 393), seed length (SL; n = 377), and seed number per panicle (SNPP; n = 376). The rice395 dataset consists of 395 diverse Asian rice samples spanning five subpopulations: indica, japonica, tropical japonica, temperate japonica, and group v, with a total of 1,536 SNP markers. The target traits for rice395 are amylose content (AC) and seed length (SL), with complete phenotypic records available for 356 and 372 accessions, respectively. The maize301 dataset was obtained from the built-in dataset in Tassel software [45]. This dataset includes 3,093 SNP markers, with the target traits being days to pollination (Dpoll; n=276), ear diameter (EarDia; n =249), and ear height (EarHT; n=279).

### Reconstructing whole-genome sequences from SNPs

To facilitate downstream genomic prediction on the basis of whole-genome features, we developed a pipeline to reconstruct sample-specific whole-genome sequences by integrating variant data from a multi-sample Variant Call Format (VCF) file onto a reference genome.

The workflow begins with essential preprocessing steps. The reference genome FASTA file is indexed via samtools (https://github.com/samtools/samtools), and the VCF file is indexed via bcftools (https://github.com/samtools/bcftools). These indexing procedures enable efficient, random-access lookups of genomic coordinates and variant records, which is critical for the performance of the subsequent steps.

Next, a list of sample identifiers is extracted from the VCF header to enable automated, batch processing of all individuals. For each sample, a custom Python script, leveraging the pysam (https://pypi.org/project/pysam/) and pyfaidx (https://pypi.org/project/pyfaidx/) libraries, generates a personalized genome sequence. The script efficiently processes variants by iterating through the VCF records for a given sample. For each variant, it substitutes the reference allele with the sample-specific allele in a mutable copy of the reference sequence.

Finally, the generated FASTA files undergo a quality control (QC) check to ensure their integrity and accuracy. This QC involves programmatic validation of the FASTA format and a verification step where a random subset of variant positions in the output sequence is cross-referenced against the input VCF to confirm correct allele incorporation. The resulting high-quality whole-genome sequences provide a reliable foundation for subsequent whole-genome feature embedding.

### Generate feature embedding for whole-genome sequences and SNP context

DNABERT-2 is a genomic foundation model based on the Transformer architecture, distinguishing itself from DNABERT through several key innovations. Unlike DNABERT, which uses k-mer tokenization, DNABERT-2 employs Byte Pair Encoding (BPE), a statistical data compression algorithm [46], for DNA sequence tokenization. Additionally, it replaces learned positional embeddings with Attention with Linear Biases (ALiBi), eliminates input length constraints [47], and integrates Flash Attention to increase computational efficiency [48]. Pretrained on datasets spanning 135 species across 7 categories, DNABERT-2 exhibits robust cross-species genomic generalization capabilities, making it a versatile tool for generating genomic feature embedding.

Owing to significant computational resource limitations, it was not feasible for us to exhaustively test all possible context lengths or a very wide spectrum of them. Therefore, we adopted an exploratory approach by evaluating a range of SNP contextual sequences, specifically from 500 bp to 3000 bp flanking the target SNPs (Figure 1A). This was intended to preliminarily investigate how varying lengths of neighboring genomic information influence trait prediction accuracy. To obtain SNP context sequences, we began by systematically identifying all heterozygous (0/1) and homozygous (1/1) SNP loci for each sample from the VCF files. For each identified mutation site, we extracted flanking sequences of varying lengths, ranging from 500 bp to 3000 bp on each side of the SNP. This resulted in total context lengths of 1001 bp to 6001 bp (assuming that the SNP itself is 1 bp). During this extraction, we ensured that these sequence fragments remained within chromosomal boundaries. To address regions with high SNP density and prevent redundant feature generation from overlapping contexts, we implemented a collision avoidance protocol: if a secondary SNP fell within a genomic region already captured as a context for a primary SNP, that secondary SNP was not processed with its own independent flanking sequences.

**Figure 1.**
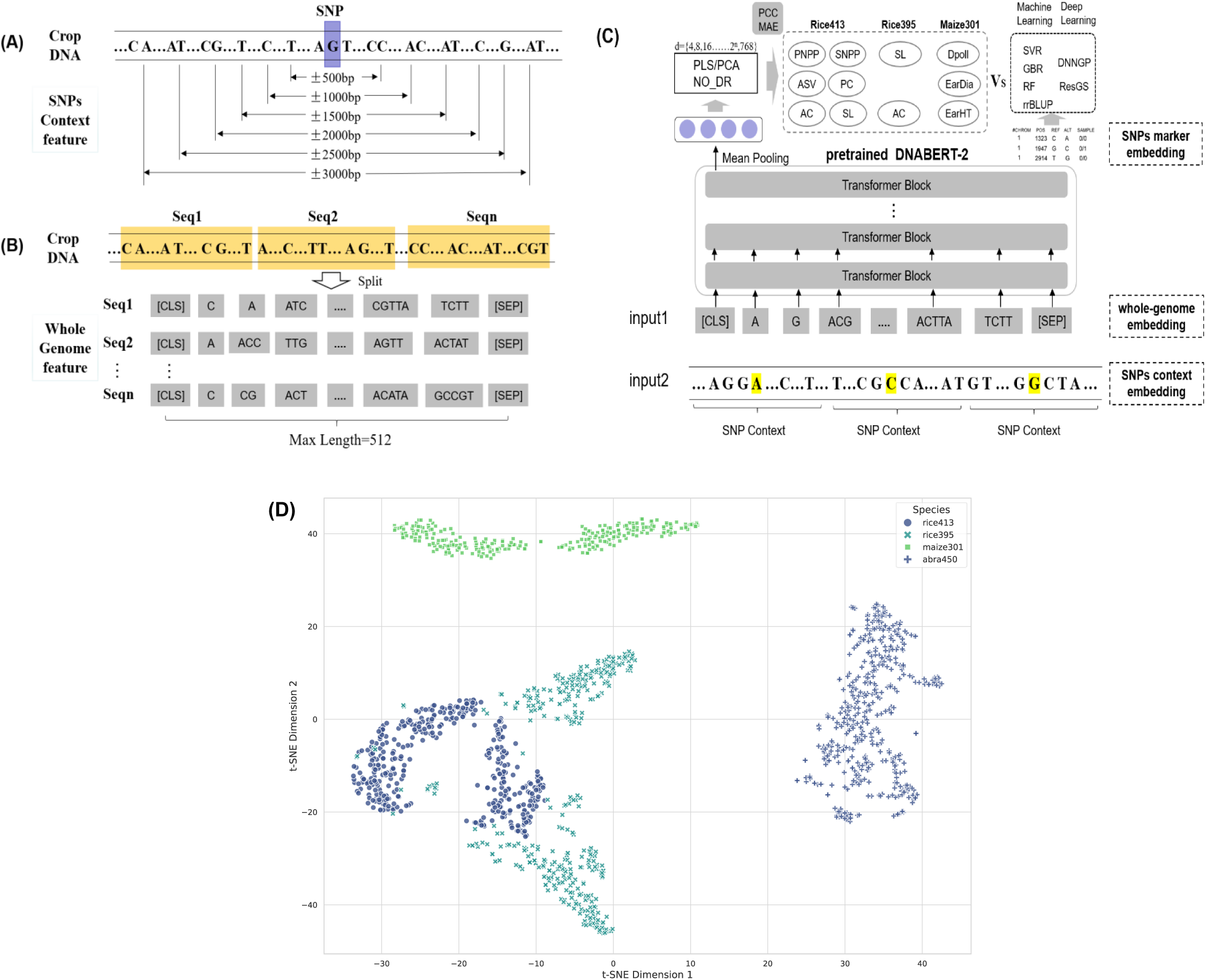
Framework for predicting crop phenotypes on the basis of two distinct embedding modes. (A) Example of the SNP context feature embedding mode. The symbol “±” denotes the extraction of a specific number of base pairs upstream and downstream from the current SNP locus. (B) Example for whole-genome feature embedding mode. The genome is divided into consecutive sections (Sec1, Sec2, …, Secn), each further partitioned into subsections with a maximum input length of 512 base pairs. (C) Generation of crop genomic feature vectors under two embedding modes via the DNABERT-2 pretrained model. (D) t-SNE visualization of high-dimensional whole-genome vectors for multiple plant species (e.g., rice, maize, Arabidopsis).

These sequences are subsequently tokenized into a sequence of tokens where frequent k-mers or subsequences are represented as individual tokens by BPE. Consequently, a 1001 bp sequence, for example, will generally be tokenized into fewer than 1001 tokens, although the exact number depends on the sequence’s composition and the BPE vocabulary. For those SNP contexts where the BPE-tokenized sequence still exceeded the maximum 512-token limit, we processed the sequence by dividing it into nonoverlapping windows of the maximum length (512 tokens). Each window was then fed through DNABert-2, and we subsequently applied average pooling across the representations derived from all these windows to generate a comprehensive embedding for the SNP context.

In parallel, we processed the reconstructed whole-genome sequences for each sample from the three crop datasets. First, we removed ambiguous bases (“N”) that can arise from sequencing artifacts. Each cleaned whole-genome sequence was subsequently partitioned into its constituent chromosomal segments. Adopting the same strategy used for long SNP contexts, we then processed each chromosomal segment. If a BPE-tokenized segment exceeded the 512-token input limit of DNABERT-2, we applied a nonoverlapping sliding window approach to divide it into 512-token chunks (Figure 1B). Each chunk was then fed into DNABERT-2 to generate a corresponding embedding. Finally, all embedding vectors produced from the chunks of a single sample’s genome were aggregated via average pooling to derive one comprehensive whole-genome embedding vector.

### Dimensionality reduction techniques

To minimize the noise unrelated to target traits in high-dimensional feature vectors, we perform dimensionality reduction on the encoded feature. Principal component analysis (PCA)—a widely used unsupervised dimensionality reduction technique—operates by transforming the original dataset’s variables into a smaller set of uncorrelated principal components via linear transformation. This process retains the most significant information from the original data while reducing dimensionality. The resulting principal components, which are linear combinations of the original features, are mutually orthogonal. By eliminating redundant information, PCA reduces noise in high-dimensional data, thereby improving the accuracy of downstream analyses [49]. This method is particularly valuable in metabolomics for analyzing high-dimensional chromatography and mass spectrometry data [50, 51].

Partial least squares (PLS), which was first introduced by Herman Wold in the 1960s [52], integrates the strengths of principal component analysis (PCA), canonical correlation analysis (CCA), and multiple regression analysis (MRA). It is particularly effective in scenarios where multicollinearity exists among independent variables. As a supervised feature extraction technique, PLS is widely used in gene expression microarray data analysis. Its primary goal is to identify orthogonal projection directions (principal components) that maximize the covariance between the projected independent variables and the dependent variable, thereby facilitating the construction of predictive models.

### Method implementation

Given the computational intensity of end-to-end fine-tuning for large models, we employed the pretrained DNABERT-2 as a powerful feature extractor to generate embeddings for our whole-genome and SNP context sequences. DNABERT-2 is a Transformer-based DNA language model built on 12 encoder layers, with a hidden size of 768, 12 attention heads, and approximately 117 million parameters (Figure 1C). The model tokenizes DNA sequences into subsequence units via BPE and processes them through its attention-driven encoder stack to learn rich genomic representations. Consistent with standard BERT architectures, it utilizes special tokens for structuring input: [CLS] is prepended to each sequence to produce an aggregate representation for sequence-level tasks; [SEP] marks the end of sequences or separates distinct segments; and [PAD] tokens are used for batch padding, which are ignored by the attention mechanism to preserve the integrity of the contextual embeddings.

We employed the non-linear dimensionality reduction technique, t-SNE, to project the high-dimensional whole-genome feature vectors from three datasets onto a two-dimensional space for visualization (Figure 1D). In the plot, each point represents a genome sample. It can be observed that Rice, Maize, and Arabidopsis (abra) form three distinct clusters, respectively. This indicates that the genomic vector embeddings generated by the DNABERT-2 model are capable of distinguishing the deep-level genomic sequence pattern differences between species. Furthermore, the two clusters belonging to the same rice species, rice413 and rice395, are closely adjacent with partial overlap. This is likely because rice395 and rice413 share the vast majority of their genomic background, yet still possess differences at key loci that determine their genetic traits. This demonstrates that the high-dimensional genomic vectors can accurately capture both the phylogenetic similarity and the subtle differences among different varieties of the same species.

Given that whole-genome embeddings may contain considerable noise or information that is not directly relevant to specific traits, we utilized dimensionality reduction techniques—such as PLS or PCA. The objective of this process was to distill trait-associated signals from the embeddings while minimizing the influence of noisy dimensions. Considering that different crop datasets contain varying numbers of valid samples (with non-missing values) across distinct phenotypes, we established a systematic search space for the optimal number of latent dimensions (d). This space was defined by an exponential series: {d | d = 2^n^, where n is an integer from 2 up to ⌊log₂N⌋}, with N being the number of samples remaining after removing those with missing phenotypic data. We then used two distinct techniques to reduce the dimensionality of the genomic embeddings: principal component analysis (PCA), an unsupervised method, and partial least squares (PLS), a supervised method that incorporates phenotype information to guide dimensionality reduction.

We conducted exhaustive hyperparameter optimization through a grid search across multiple machine learning architectures. For benchmarking, we implemented several well-established genomic prediction methods, including support vector regression (SVR), ridge regression best linear unbiased prediction (rrBLUP), random forest (RF), and gradient boosting regression (GBR), by leveraging the automated machine learning framework provided by PyCaret (https://pycaret.org/). This platform streamlined comparative analysis by handling data preprocessing, feature engineering, and model selection through its integrated pipeline.

### Indicator evaluation and calculation

To ensure robust performance estimation, we implemented tenfold cross-validation across all datasets. This validation approach partitions the data into ten subsets, with nine for training and one for testing in each iteration. We report the mean PCC values across all folds as our primary accuracy metric, providing a comprehensive assessment of model generalizability.

As a complementary measure of prediction accuracy, we calculated the MAE for quantitative trait predictions. This metric provides an intuitive measure of average prediction error magnitude in the original units of measurement, offering additional insights into model performance beyond correlation-based metrics.

The mathematical formulations for these evaluation metrics are provided in Equations (1) - (2), with detailed derivations available in the Supplementary Materials. This dual-metric approach ensures comprehensive assessment of both the strength of association (PCC) and absolute prediction accuracy (MAE) across all tested phenotypes.

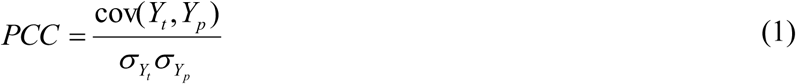

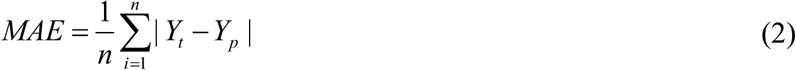

where cov represents the covariance between the true phenotypic values *Y*_t_ and the predicted phenotypic values *Y_p_*, which represent the standard deviations of the true phenotypic values *Y*_t_ and the predicted phenotypic values *Y_p_*, respectively.

We assessed trait heritability via broad-sense heritability (h²) following established genomic prediction protocols [35]. This comprehensive metric quantifies the proportion of total phenotypic variance attributable to all genetic components, including additive effects, dominance variance, and epistatic interactions across the rice413, rice395, and maize301 populations.

The broad-sense heritability was calculated as follows:

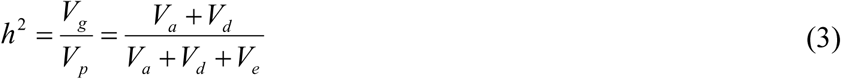

This formulation enables a robust assessment of the genetic contribution to phenotypic variation, accounting for both genetic and environmental influences. In the formula, *V_a_* denotes the additive genetic variance, *V_d_* denotes the dominant genetic variance, and *V_e_* denotes the environmental variance. In our study, variance components (*V_a_*, *V_d_*, *V_e_*) were derived through genome-based restricted maximum likelihood (GREML) analysis, a well-established method in quantitative genetics. The core approach leverages a mixed linear model framework where the genomic relationship matrix (GRM) model genetic similarity among individuals on the basis of genome-wide SNPs, followed by variance component decomposition via restricted maximum likelihood (REML). This methodology specifically partitions the observed phenotypic variance into additive genetic variance (*V_a_*) and environmental variance (*V_e_*), with narrow-sense heritability calculated as h² = *V_a_* / (*V_a_* + *V_e_*). Notably, our implementation focuses on additive effects (*V_a_*) as the primary genetic architecture for the studied traits, which is consistent with common practices in plant genomic studies, whereas dominance variance (*V_d_*) was not estimated because both sample size considerations and prior evidence suggest minimal dominance effects for these traits.

## Results

### Phenotypic prediction results under the SNP-context feature embedding mode

We adapted contextual analysis approaches from natural language processing to investigate how genetic information in SNP-flanking regions influences phenotypic prediction accuracy. For each SNP variant, we extracted genomic segments of varying lengths (500–3000 bp) centered on the polymorphic site. These “context windows,” which included the variant itself, were then processed through our feature embedding pipeline to capture local genomic information. To assess prediction performance, we implemented four representative models: SVR, rrBLUP, RF, and GBR. Finally, we evaluated the ability of each model to predict phenotypic traits in the rice413 dataset via features derived from the different context window sizes.

In Figure 2, the horizontal axis labels (±500 to ±3000) indicate the length (in bp) of the genomic sequence extracted upstream and downstream of each SNP variant. The coordinate at ‘0’ represents the baseline prediction method using only SNP markers, with its PCC value taken from the literature [32]. The yellow circles highlight the highest PCC achieved for each trait via our contextual models across the tested window sizes (500 bp to 3,000 bp).

**Figure 2.**
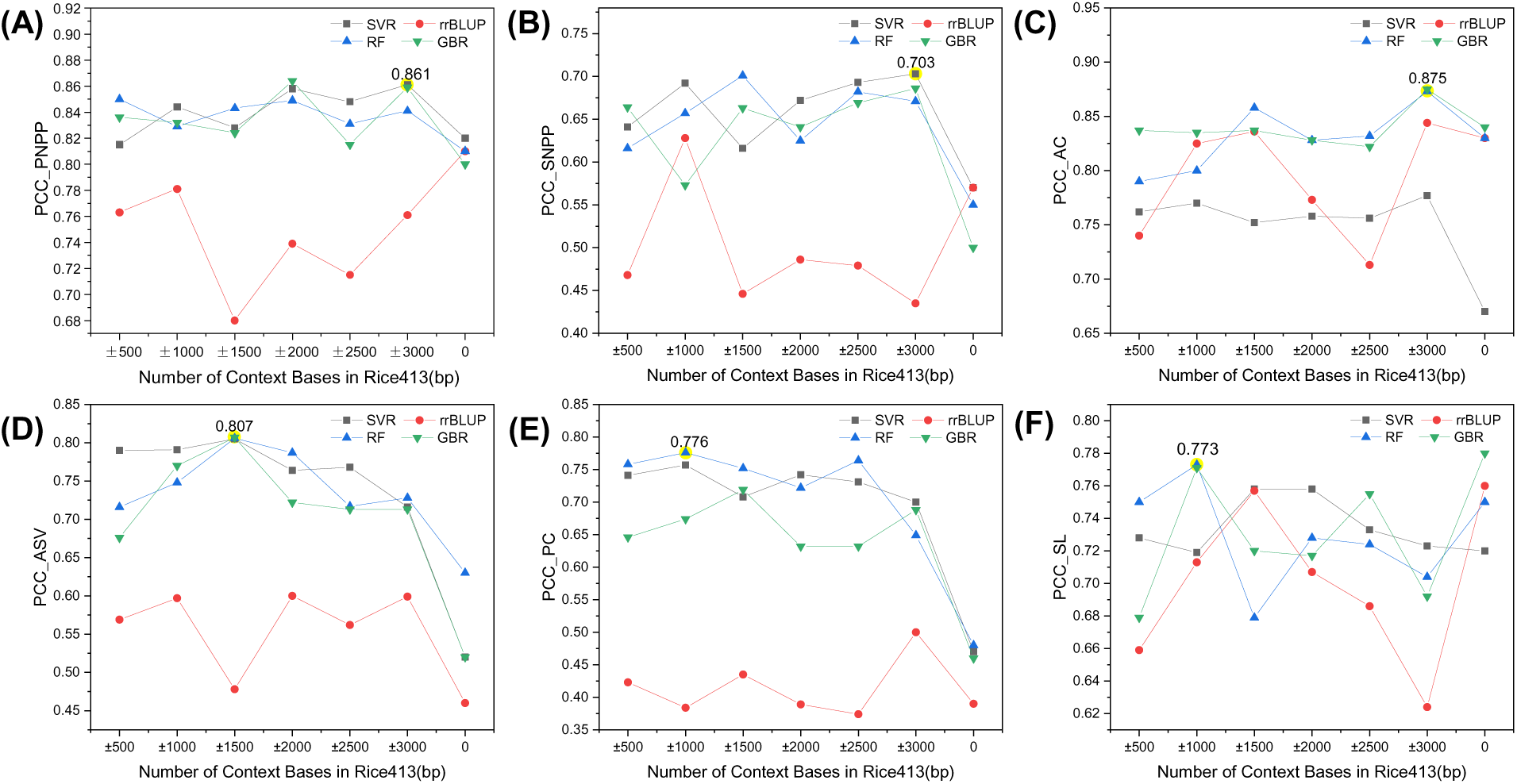
Phenotypic prediction accuracy for six agronomic traits in rice413 via SNP-context features (500 bp--3000 bp) across four ML algorithms. (A) PNPP; (B) SNPP; (C) AC; (D) ASV; (E) PC; (F) SL. Prediction performance was evaluated by the PCC for the SVR, rrBLUP, RF, and GBR models.

The optimal prediction accuracies for specific traits were achieved at different context lengths: 3,000 bp for PNPP, SNPP, and AC; 1,500 bp for ASV; and 1,000 bp for PC. Notably, the PCC between the predicted and true phenotypic values was generally greater than that of the SNP-only baseline. Fȯr the SL trait, some models performed better than or comparably to the baseline when context lengths between 1,000 and 2,000 bp were used. These findings underscore the importance of incorporating flanking genomic sequences to increase the accuracy of phenotypic prediction.

For traits such as PNPP, SNPP, and PC, rrBLUP achieves lower prediction accuracy than nonlinear models (e.g., SVR and GBR) when contextual genomic features are used. However, compared with SNP-context embeddings, rrBLUP performs better for seed length (SL) when raw SNP markers are utilized. We speculate that these observed differences stem from how these algorithms process genetic information. As a linear mixed model, rrBLUP is optimized for capturing additive genetic effects and relies on the direct estimation of allele dosage effects from discrete SNP markers. For SL, which is often highly polygenic and where additive variance plays a dominant role, the transformation of these discrete SNPs into continuous, dense embedding vectors might inadvertently obscure or dilute these direct additive signals. The embedding process, while capturing the local context, might make it more challenging for a linear model (rrBLUP) to precisely quantify the individual additive contributions that it excels at modeling. Conversely, for other traits, such as PNPP, SNPP, AC, ASV, and PC, nonlinear algorithms (e.g., SVR and GBR) demonstrate the ability to benefit from SNP-context embeddings. This is likely because these models are inherently capable of capturing more complex, nonlinear relationships and feature interactions. The embeddings likely provide a richer feature representation for these traits by encapsulating higher-order sequence patterns, local contextual information, and potential proxies for epistatic or context-dependent genetic effects. SVR and GBR are well suited for exploiting these complex patterns, which may be more characteristic of the genetic architecture of traits such as PNPP, SNPP, and PC, leading to improved predictive performance.

As shown in Figure 3, for the AC trait in the rice395 dataset, SVR significantly outperformed the other algorithms across all tested context lengths (500–3000 bp). It achieved its peak predictive performance (PCC = 0.609) at the 3,000 bp context length. For the SL trait, SVR, RF, and GBR also outperformed rrBLUP when SNP-context features were used, with RF attaining the highest accuracy (PCC = 0.482) at a 1,500 bp context length. We can find that both the AC and SL traits in this dataset is that prediction accuracies using SNP-context features were lower than those achieved with traditional SNP markers alone.

**Figure 3.**
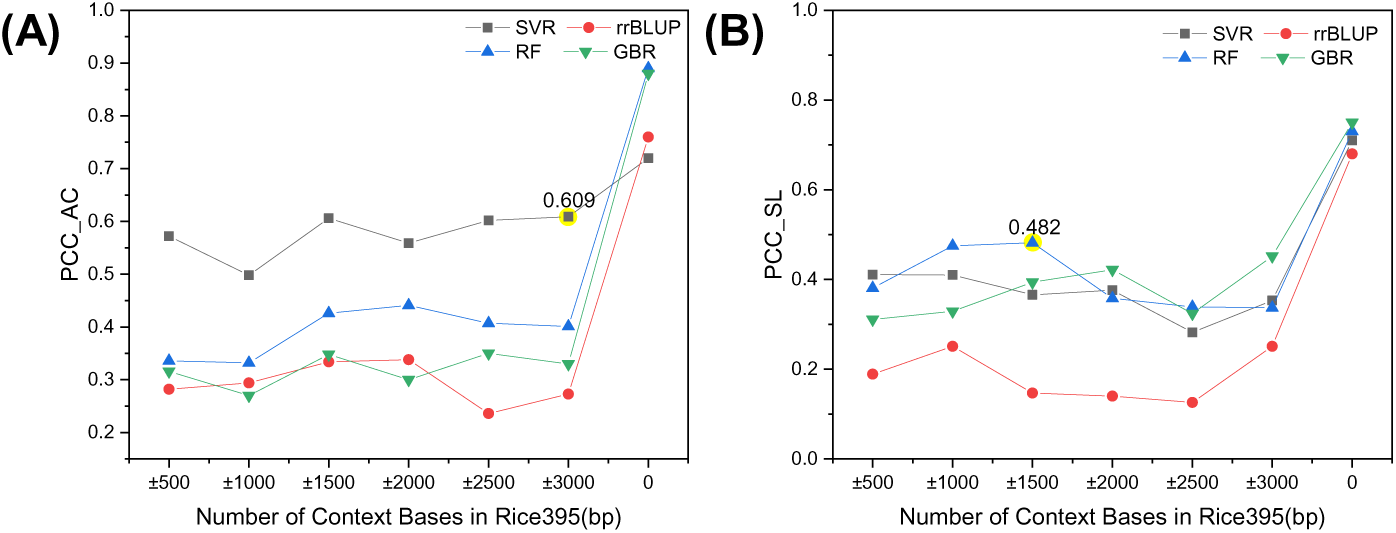
Prediction accuracy for AC and SL traits in rice395 via four ML algorithms with SNP-context features (±500 bp to ±3000 bp). (A) AC; (B) SL. Performance was evaluated by the PCC for the SVR, rrBLUP, RF, and GBR models.

Figure 4 show that prediction accuracy using SNP-context features varied significantly among algorithms. Notably, several configurations outperform traditional SNP marker-based approaches. SVR demonstrates the strongest performance for Dpoll prediction, achieving a peak accuracy of 0.742 at a context length of 3000 bp.

**Figure 4.**
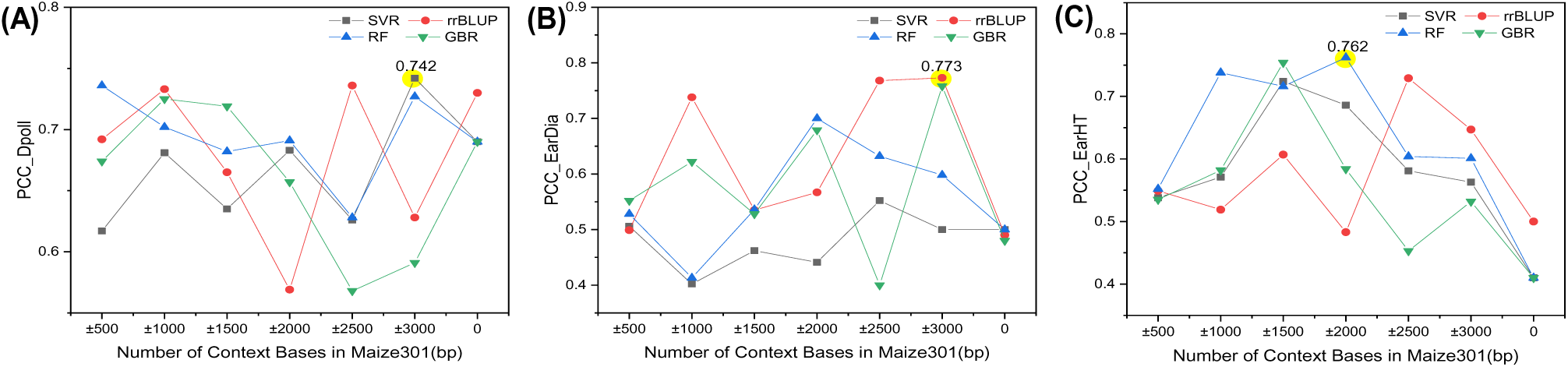
Prediction accuracy in maize301 using SNP-context features (500--3000 bp) across four machine learning algorithms. (A) Days to pollination (Dpoll); (B) Ear diameter (EarDia); (C) Ear height (EarHT). Performance was evaluated via the PCC.

For EarDia, rrBLUP shows optimal performance, with an accuracy of 0.773 at 3000 bp. RF performs best for EarHT prediction, reaching 0.762 accuracy at a 2000 bp context length. These results highlight the context length-dependent nature of prediction performance across different traits.

Comparative evaluation demonstrates consistent improvements when using SNP-context features compared to traditional SNP markers. The average accuracy gains are most substantial for EarHT (35.2%), followed by EarDia (28.3%) and Dpoll (5.2%). This pattern suggests that the relative benefit of contextual sequence information varies considerably among traits, potentially reflecting differences in their underlying genetic architectures.

### Phenotypic prediction results based on whole-genome feature embedding

The genetic regulation of crop traits involves complex interactions among multiple major genes, minor genes, and quantitative trait loci (QTLs) distributed throughout the genome. These genetic elements often span multiple chromosomes and may exhibit nonlinear epistatic interactions, presenting significant challenges for prediction methods that rely solely on SNPs or localized sequence contexts.

To overcome these limitations, we implemented a whole-genome feature embedding strategy that captures nucleotide sequence information across complete chromosomal sequences. This comprehensive approach was complemented by dimensionality reduction through both PCA and PLS. By systematically evaluating prediction performance across different reduced dimensions, our framework provides an enhanced capacity to model the distributed genetic architecture underlying complex crop traits.

Our comprehensive evaluation reveals significant variation in prediction accuracy across different ML algorithms and feature dimensionality approaches (Figures 5 and S1). The results demonstrate clear trait-specific patterns in optimal prediction strategies.

**Figure 5.**
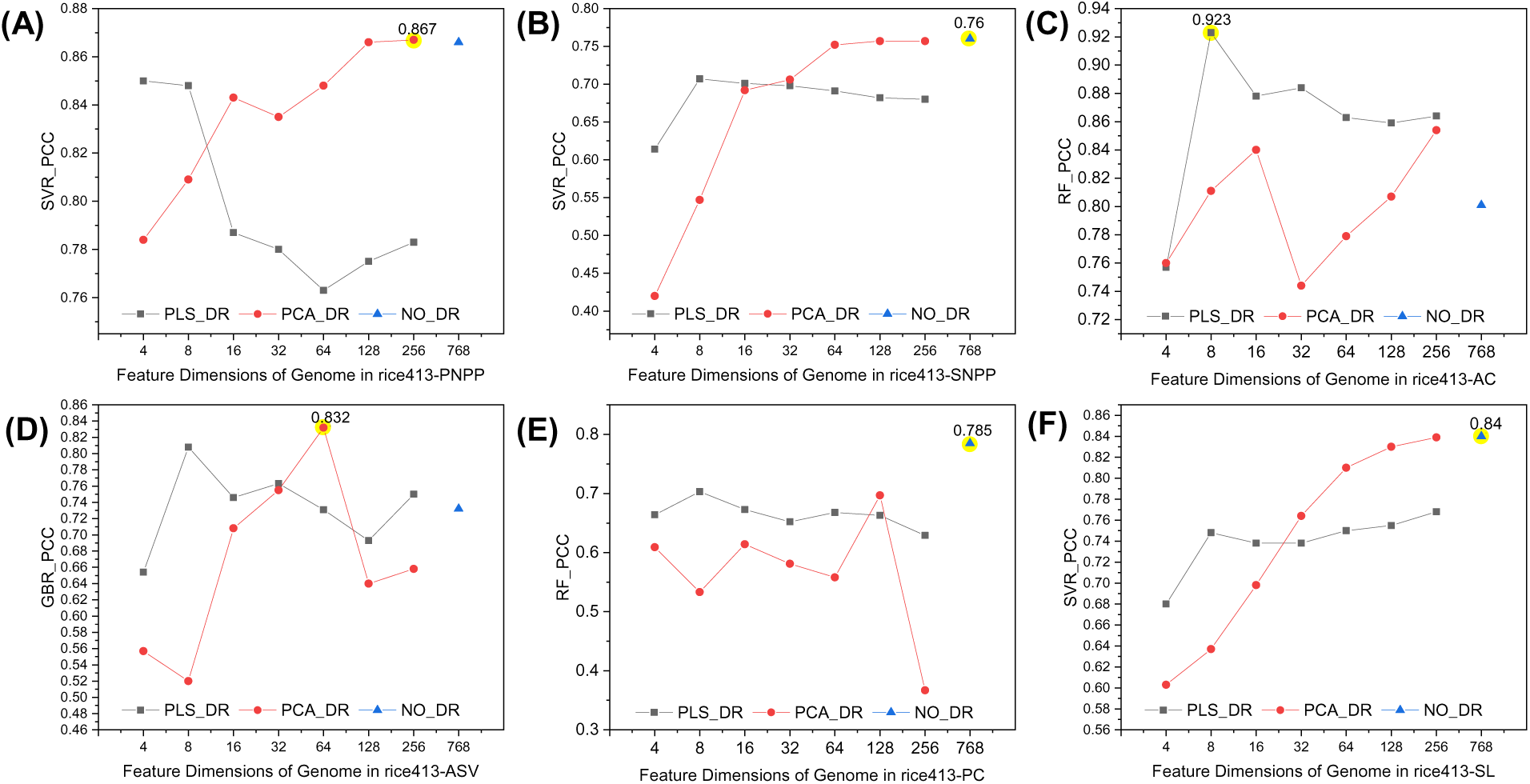
Comparative performance of whole-genome feature embedding with dimensionality reduction (PLS/PCA) versus full-dimensional (768D) approaches for six agronomic traits in rice413. (A) PNPP; (B) SNPP; (C) AC; (D) ASV; (E) PC; (F) SL. The yellow highlights denote the optimal performance among the four ML algorithms (SVR, rrBLUP, RF, and GBR). The complete algorithm comparisons are shown in Figure S1.

For PNPP, PCA demonstrates superior performance to PLS at higher dimensionalities (>16 components). SVR achieves maximal prediction accuracy (PCC = 0.867) at 256 dimensions, matching the performance of full-dimensional embedding. Similar optimal performance is observed for SNPP, where SVR with complete feature sets yields the highest accuracy (PCC = 0.76). Interestingly, for PNPP, the PLS performance decreases, whereas the PCA performance improves as the target dimensionality increases. This phenomenon likely stems from fundamental operational differences between the methods and the nature of the PNPP’s genetic signal within the high-dimensional embedding space. As a supervised method, PLS aims to find components that maximize covariance between the 768-dimensional embeddings and the PNPP trait. When the number of PLS components increases (from 4 to 256), it attempts to explain more PNPP variance by incorporating additional, potentially less robust, relationships from the original embeddings. However, these supplementary PLS components might increasingly capture noise or spurious correlations from the training data rather than robust predictive signals. Conversely, PCA, an unsupervised method, identifies principal components based solely on maximum variance within the embedding data, without regard to the PNPP trait. If the PNPP’s genetic architecture involves numerous, subtle signals dispersed throughout the high-dimensional embedding space, retaining more PCA components captures more of this total variance and, consequently, more of these diffuse, trait-relevant signals.

Notably, distinct patterns emerge for quality-related traits. AC shows markedly improved prediction via PLS dimensionality reduction, particularly when combined with RF, which reaches peak accuracy (PCC = 0.923) at 8 dimensions. For ASV, GBR with PCA at intermediate dimensionality (64 components) provides optimal results (PCC = 0.832). PC and SL both achieve maximum accuracy without dimensionality reduction when RF (PCC = 0.785) and SVR (PCC = 0.84) are used, respectively.

As shown in Figure 6 and S2 (a-f), the prediction accuracy for the AC trait in the rice395 dataset remains relatively stable with increasing dimensionality when SVR is used, which demonstrates superior consistency over other algorithms. The highest prediction accuracy for AC (PCC=0.561) is achieved when GBR is combined with PCA at 64 dimensions. For the SL trait, the highest prediction accuracy (PCC=0.5) is achieved via GBR with PLS at 64 dimensions. These results suggest that dimensionality reduction techniques, particularly at intermediate dimensions (e.g., 64), can increase the prediction accuracy for specific traits. However, the performance varies significantly across traits and algorithms, highlighting the importance of tailoring dimensionality reduction and algorithm selection to the specific trait of interest. Additionally, the relatively lower prediction accuracy for SL compared to AC may reflect the greater genetic complexity or environmental sensitivity.

**Figure 6.**
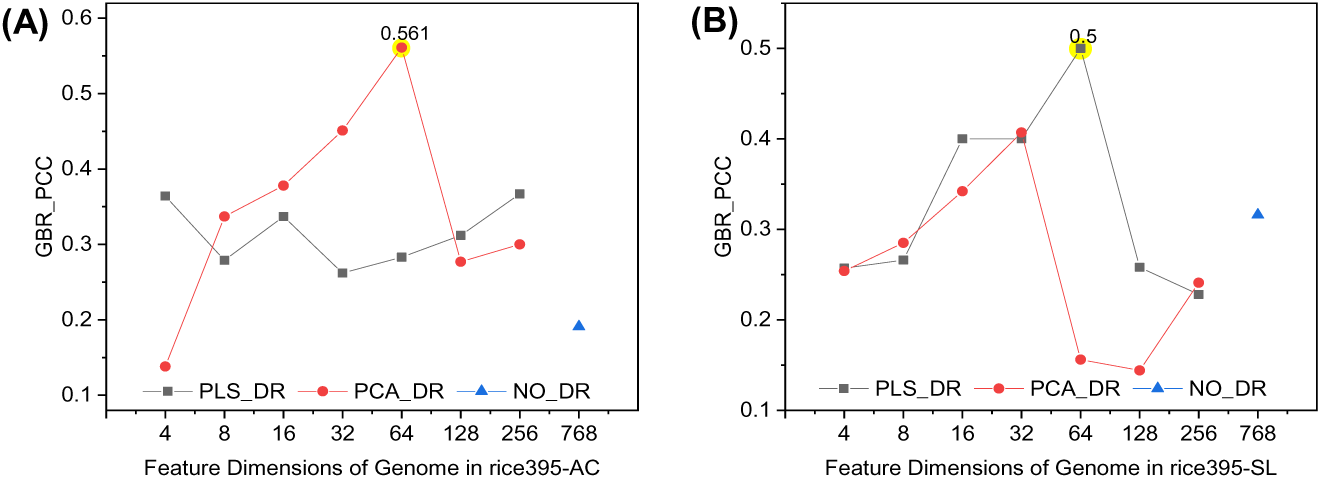
Prediction accuracy for AC and SL traits in rice395 via whole-genome feature embedding with dimensionality reduction (PLS, PCA) or full dimensions (768). (A) AC; (B) SL. Yellow highlights indicate the highest accuracy among the four algorithms (SVR, rrBLUP, RF, and GBR). The complete algorithm comparisons are shown in Figure S2.

We observe that PLS dimensionality reduction outperforms PCA when SVR is used to predict Dpoll, EarDia, and EarHT (Figure 7). For Dpoll, the highest prediction accuracy (PCC=0.806) is achieved via rrBLUP with PCA at 16 dimensions, representing a 7.6% improvement over traditional SNP feature embedding. For EarDia, GBR without dimensionality reduction yields the highest accuracy (PCC=0.849), a 35.9% improvement over traditional methods. For EarHT, rrBLUP combined with PCA at 128 dimensions achieves the maximum accuracy (PCC=0.746), a 24.6% increase over traditional SNP feature embedding.

**Figure 7.**
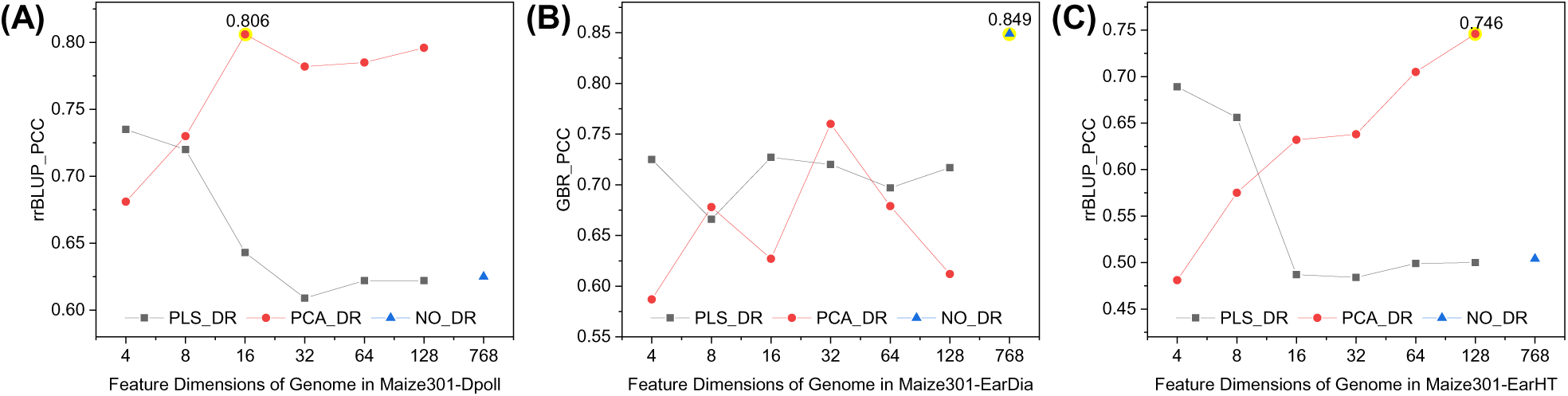
Prediction accuracy for maize301 traits via whole-genome feature embedding with dimensionality reduction (PLS, PCA) or full dimensions (768D). (A) Dpoll; (B) EarDia; (C) EarHT. Yellow highlights indicate optimal performance among the four algorithms (SVR, rrBLUP, RF, and GBR). The complete results are shown in Figure S3.

The highest average prediction accuracies for each trait across the rice413, rice395, and maize301 datasets using whole-genome feature embedding, SNP context feature embedding, and traditional SNP features (“SNP*” column) are summarized in Table 1∼3. The “SNP*” column represents the optimal traditional ML algorithm based on SNP marker features, with its values derived from the literature [32]. When the best-performing algorithms differ between SNP context embedding and whole-genome embedding, the SNP* column reflects the highest precision value from either algorithm. Our results show that no single algorithm consistently achieves the best prediction performance across all traits, which is consistent with findings from prior studies [53, 54]. Furthermore, certain traits exhibit higher prediction accuracy when whole-genome feature embedding is combined with PCA or PLS dimensionality reduction, compared to using the full-dimensional embeddings.

**Table 1.**
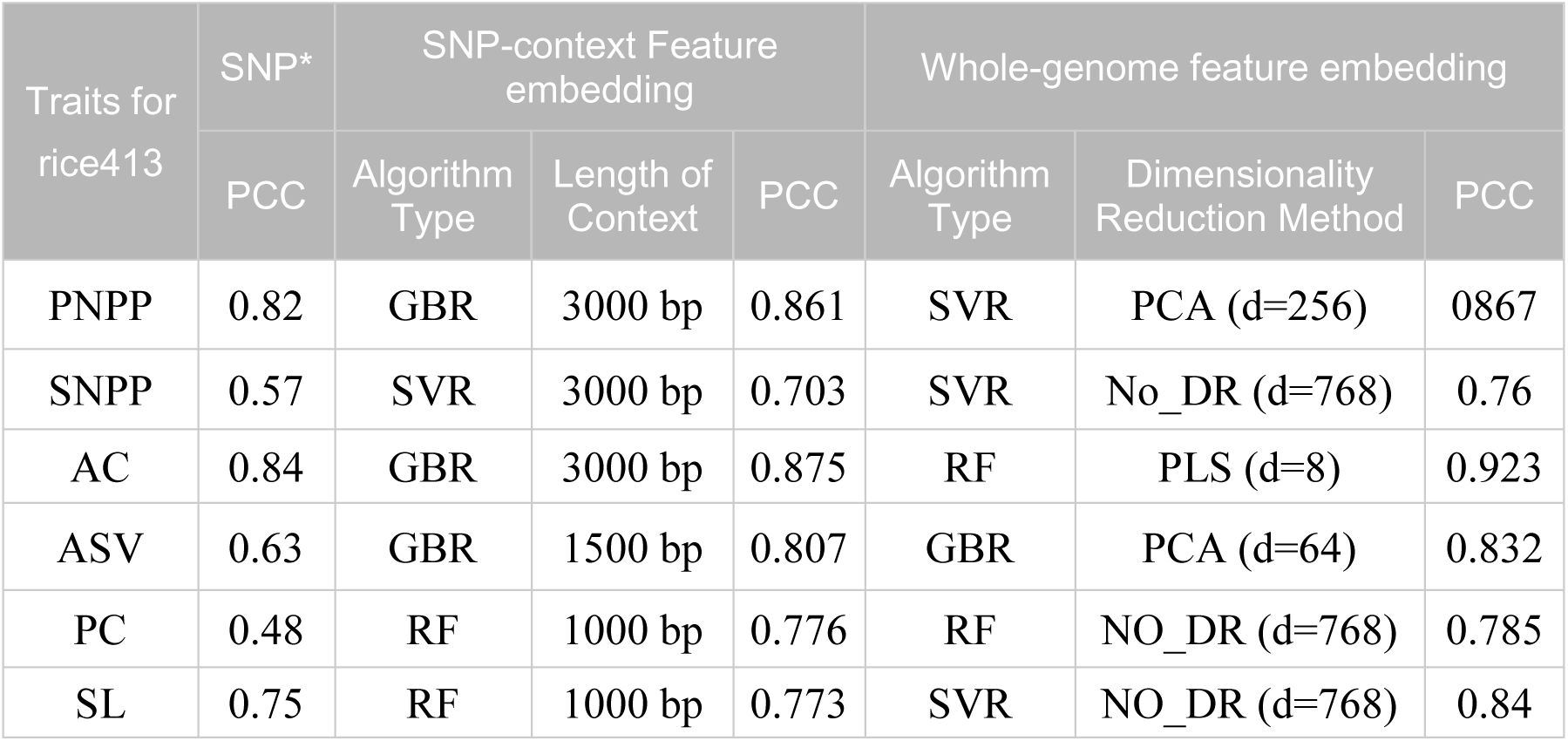
Highest average prediction accuracy for various traits in rice413 under different feature modes.

| Traits for rice413 | SNP* | SNP-context Feature embedding |  |  | Whole-genome feature embedding |  |  |
| --- | --- | --- | --- | --- | --- | --- | --- |
|  | PCC | Algorithm Type | Length of Context | PCC | Algorithm Type | Dimensionality Reduction Method | PCC |
| PNPP | 0.82 | GBR | 3000 bp | 0.861 | SVR | PCA (d=256) | 0.867 |
| SNPP | 0.57 | SVR | 3000 bp | 0.703 | SVR | No_DR (d=768) | 0.76 |
| AC | 0.84 | GBR | 3000 bp | 0.875 | RF | PLS (d=8) | 0.923 |
| ASV | 0.63 | GBR | 1500 bp | 0.807 | GBR | PCA (d=64) | 0.832 |
| PC | 0.48 | RF | 1000 bp | 0.776 | RF | NO_DR (d=768) | 0.785 |
| SL | 0.75 | RF | 1000 bp | 0.773 | SVR | NO_DR (d=768) | 0.84 |

Figure 8 illustrates the PCC values for all traits in the rice413, comparing the performance of the optimal machine learning (ML) model under three different input feature types—traditional SNP markers, SNP-context features, and whole-genome features—against that of state-of-the-art deep learning models which use SNP markers as input. For the PNPP trait, machine learning models using whole-genome feature embedding demonstrates predictive performance nearly equivalent to that of SNP-context feature embedding, with marginal accuracy improvements of 2.7% and 0.7% over those of the DNNGP and ResGS models based on SNP markers, respectively. Notably, the SNPP trait exhibits superior prediction accuracy when whole-genome feature embedding is utilized, outperforming SNP-context feature embedding by 5.7% and substantially exceeding the DNNGP and ResGS models by 10% and 9%.

**Figure 8.**
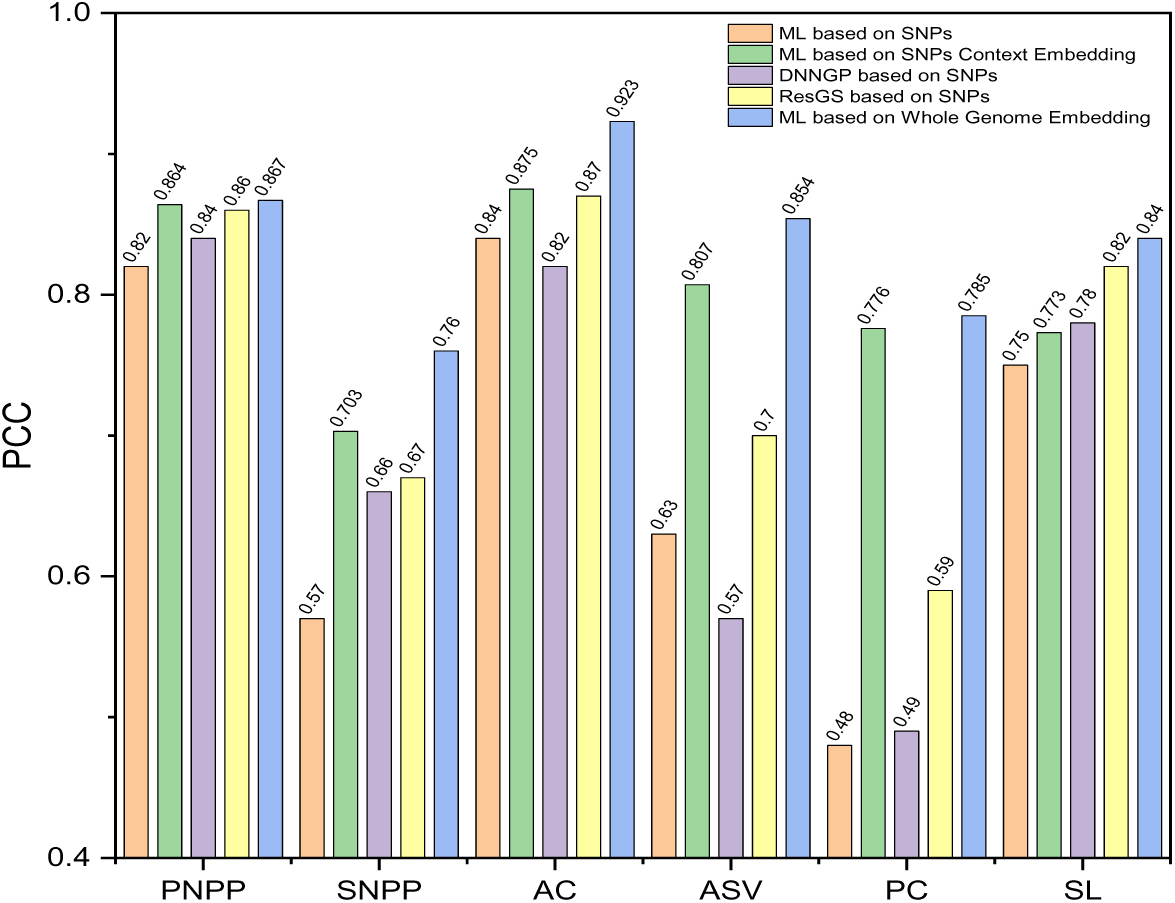
Comparison of prediction accuracy among traditional ML algorithms, which represent the type of algorithm that achieves optimal performance for each trait in Table 1 under different feature modes. DNNGP and ResGS are based on SNP markers, and ML algorithms are based on SNP context feature embedding and whole-genome feature embedding for PNPP, SNPP, AC, ASV, PC, and SL in Rice413.

With respect to the AC trait, ML based on whole-genome feature embedding achieves a 4.8% greater accuracy than SNP-context feature embedding but outperforms the DNNGP and ResGS models by 10.3% and 5.3%. The most remarkable performance enhancement is observed in the ASV trait, where ML based on whole-genome feature embedding demonstrates a 4.7% improvement over SNP-context feature embedding and substantial advantages of 28.4% and 15.4% over the DNNGP and ResGS models.

For the PC trait, models using whole-genome feature embedding perform slightly better than those that use SNP-context features. However, they demonstrate significant accuracy improvements of 30.5%, 29.5%, and 19.5% when compared to the baseline methods, namely the traditional SNP-based machine learning model, DNNGP, and ResGS, respectively. For the SL trait, whole-genome feature embedding outperforms the traditional SNPs feature embedding method by 9%, and exceeds DNNGP and ResGS models by 6% and 2%.

These comparative results consistently demonstrate the superior predictive ability of whole-genome feature embedding across multiple traits. This superiority is particularly evident in its substantial performance advantages over both the traditional machine learning model using SNP features and state-of-the-art deep learning models such as DNNGP and ResGS.

Consistent with previous studies [55–58], our findings indirectly reflect that trait heritability (Table 4) significantly influences the performance of genomic prediction algorithms. The results show distinct patterns in prediction accuracy across different traits and methods. For SNPP, ASV, and PC traits, SNP-context feature embedding at specific context lengths (3000 bp, 1500 bp, and 1000 bp, respectively) substantially outperformed traditional SNP-based methods. Whole-genome feature embedding yielded further accuracy improvements of 5.7%, 4.7%, and 0.9% for these traits, respectively. This enhancement can be attributed to the incorporation of linkage disequilibrium (LD) and genetic associations between SNPs, which are typically neglected in conventional single-nucleotide polymorphism (SNP) feature modeling. The superior performance of whole-genome feature embedding may reflect DNABERT-2’s capacity to identify major genes or key QTL characteristics within extended genomic contexts. While our investigation was limited to six context lengths within a 3000 bp range, the potential for improved accuracy through longer context lengths remains unexplored. Notably, the PNPP trait showed comparable prediction accuracy across SNP-context embedding, whole-genome embedding, and the ResGS model. This is likely because that the selected SNP markers may encompass the core quantitative trait loci crucial for PNPP expression, rendering additional genomic context information redundant. Phenotypes with moderate heritability are particularly susceptible to intricate gene regulatory networks and environmental perturbations, which complicates the disentanglement of causal genetic variants from stochastic noise.

For the AC and SL traits, advanced embedding methods led to substantial accuracy improvements. SNP-context feature embedding enhanced the prediction accuracy by 3.5% and 2.3%, respectively, while whole-genome feature embedding achieved more significant improvements of 8.3% and 9%. The superior performance of whole-genome feature embedding for these traits may align with their higher heritability values, suggesting greater genetic determination. This phenomenon may reflect the complex regulation of AC and SL traits by multiple genes or QTLs, where comprehensive genomic information captures additional nonadditive and recessive effects. These findings corroborate previous observations in animal genomic prediction studies [59, 60] regarding the positive correlation between heritability and prediction accuracy.

Taken together, these results highlight the necessity of selecting feature embedding methods based on trait-specific genetic architectures. Whole-genome feature embedding emerges as a particularly powerful tool for traits with complex genetic regulation and high heritability, while context-specific embeddings may suffice for traits dominated by a few major QTLs.

The PCC primarily assesses the linear association strength between predicted and actual trait values, whereas the MAE quantifies the magnitude of deviation between the two. In this study, we also computed the MAEs for all the prediction methods applied to the rice413 dataset (Figure 9). The benchmark MAE values for traditional SNP-based prediction were sourced from the literature [32]. In instances where the algorithms achieving the highest average prediction accuracy differed between SNP context embedding and whole-genome embedding, the minimum MAE based on SNP marker features from both algorithms was adopted.

**Figure 9.**
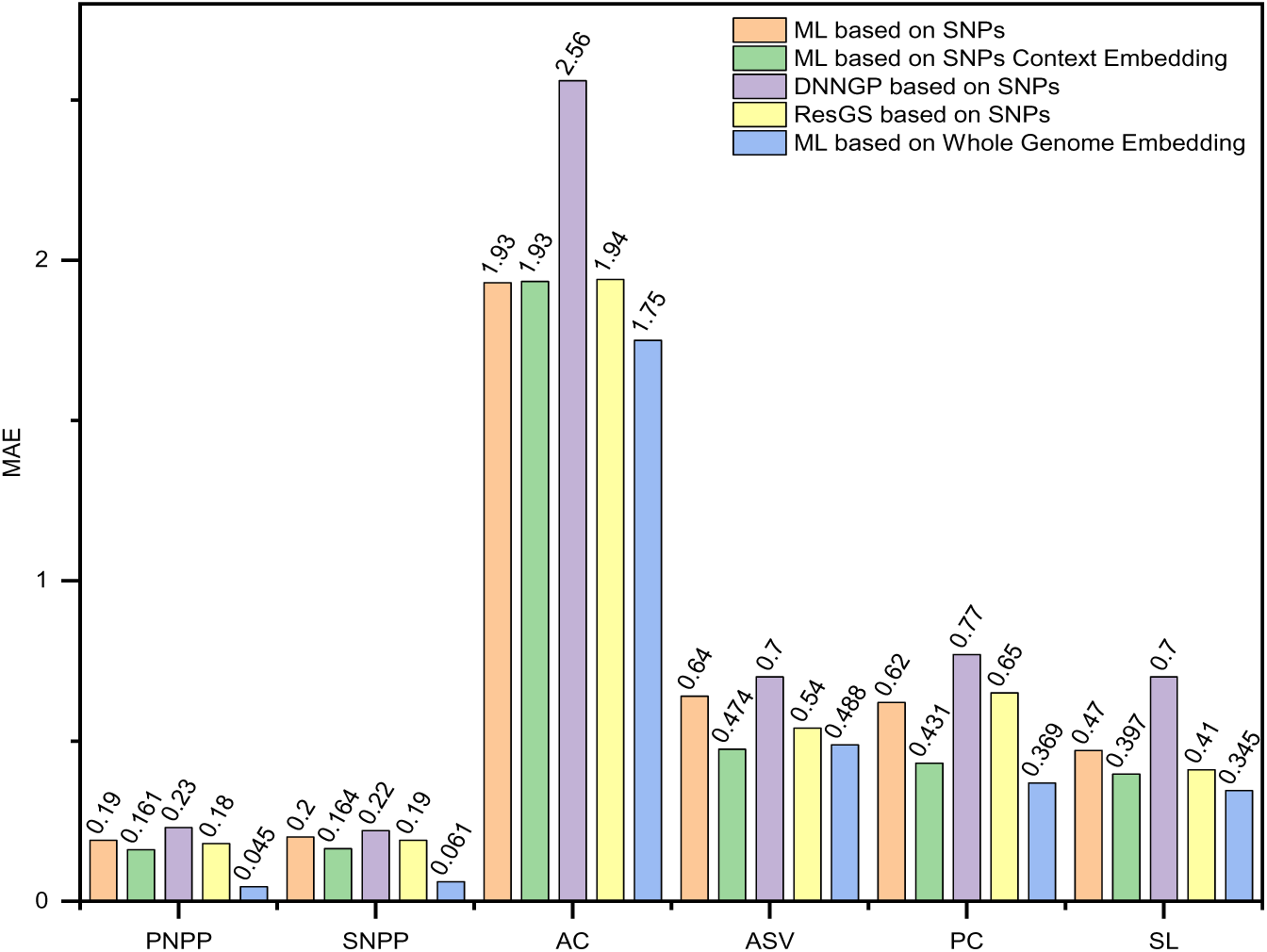
Comparison of MAEs among traditional ML algorithms, DNNGP, ResGS (based on SNP markers) and ML based on SNP context feature embedding and whole-genome feature embedding for PNPP, SNPP, AC, ASV, PC, and SL in Rice413.

Overall, ML based on whole-genome feature embedding achieved the lowest MAE for the PNPP, SNPP, AC, PC, and SL traits in the rice413 dataset. For the AC trait, the MAE from ML based on SNP-context embedding was comparable to that of traditional SNP features and the ResGS model; however, whole-genome encoding resulted in a substantially lower MAE. Similarly, the SL trait also exhibited a marked reduction in MAE when whole-genome feature embedding was used compared with the other approaches.

For traits such as ASV and PC, using SNP-context features led to a significant reduction in the MAE compared with traditional SNP methods, DNNGP, and ResGS. This improvement may be attributed to the fact that traditional SNP features typically concentrate solely on the effects of individual SNPs, often overlooking the genetic context of neighboring regions, which can result in the loss or distortion of trait‒genetic association signals. In contrast, by encoding broader genomic regions, both SNP-context and whole-genome methods can capture important long-range dependencies and distant genetic associations.

Figure 10 compares the performance of SNP-context feature embedding and whole-genome feature embedding against traditional SNP features, DNNGP, and ResGS for predicting AC and SL traits in the rice395 dataset. In contrast to the results in rice413, both novel embedding methods underperformed compared with the benchmark models, exhibiting lower PCCs and higher MAEs. However, SNP-context feature embedding achieved a 4.8% greater prediction accuracy for the AC trait than did whole-genome feature embedding, whereas whole-genome feature embedding showed marginally better performance for the SL trait.

**Figure 10.**
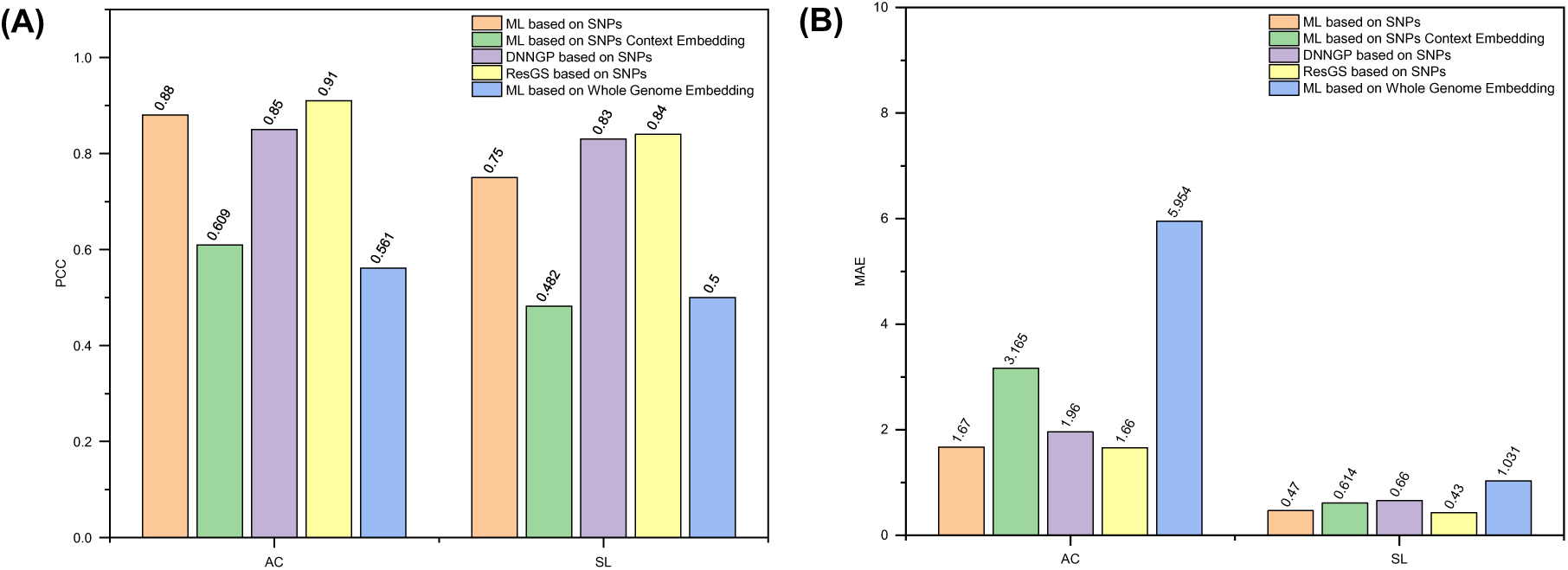
Performance comparison of traditional ML algorithms, which represent the type of algorithm that achieves optimal performance for each trait in Table 2 under different feature modes. DNNGP and ResGS are based on SNP markers, whereas ML algorithms are based on the SNP context and whole-genome feature embedding for traits in rice395. (A) PCC values for predicting AC and SL traits. (B) MAEs for predicting AC and SL traits.

**Table 2.** Highest average prediction accuracy for various traits in rice395 under different feature modes.

| Traits for<br>rice395 | SNP* | SNP-context Feature<br>embedding |  |  | Whole-genome feature embedding |  |  |
| --- | --- | --- | --- | --- | --- | --- | --- |
|  | PCC | Algorithm<br>Type | Length of<br>Context | PCC | Algorithm<br>Type | Dimensionality<br>Reduction Method | PCC |
| AC | 0.88 | SVR | 3000 bp | 0.609 | GBR | PCA (d=64) | 0.561 |
| SL | 0.75 | RF | 1500 bp | 0.482 | GBR | PLS (d=64) | 0.5 |

Further analysis revealed that the heritability of the AC and SL traits was significantly lower in rice395 than in rice413. This discrepancy may arise from differences in the number and effect sizes of genes controlling AC and SL, as well as variations in gene‒environment interactions across rice varieties. The ultra-low heritability in rice395 suggests reduced genetic stability across generations, making these traits more susceptible to environmental influences and other confounding factors. Consequently, the predictive power of SNP-context feature embedding and whole-genome feature embedding was limited, resulting in lower accuracy and higher MAE values. This observation aligns with findings from Habier *et al*. (2011), who reported that traits with lower heritability exhibit weaker gene‒trait correlations, leading to reduced accuracy in estimating marker effects across the genome.

Figure 11 compares the prediction performance of our proposed embedding methods against traditional SNP features, DNNGP, and ResGS for three traits (Dpoll, EarDia, and EarHT) in the maize301 dataset. For the Dpoll trait, whole-genome feature embedding achieved the highest prediction accuracy (PCC = 0.806) and the lowest MAE, representing improvements of 7.6%, 2.6%, and 2.6% over traditional SNP features, DNNGP, and ResGS, respectively. This method was also superior for the EarDia trait, with a prediction accuracy of 0.849, which surpassed those of the three benchmark models by 35.9%, 22.9%, and 19.9%, respectively. For the EarHT trait, SNP-context feature embedding was the top performer, outperforming traditional SNP features, DNNGP, and ResGS, with accuracy improvements of 26.2%, 19.2%, and 20.2%, respectively.

**Figure 11.**
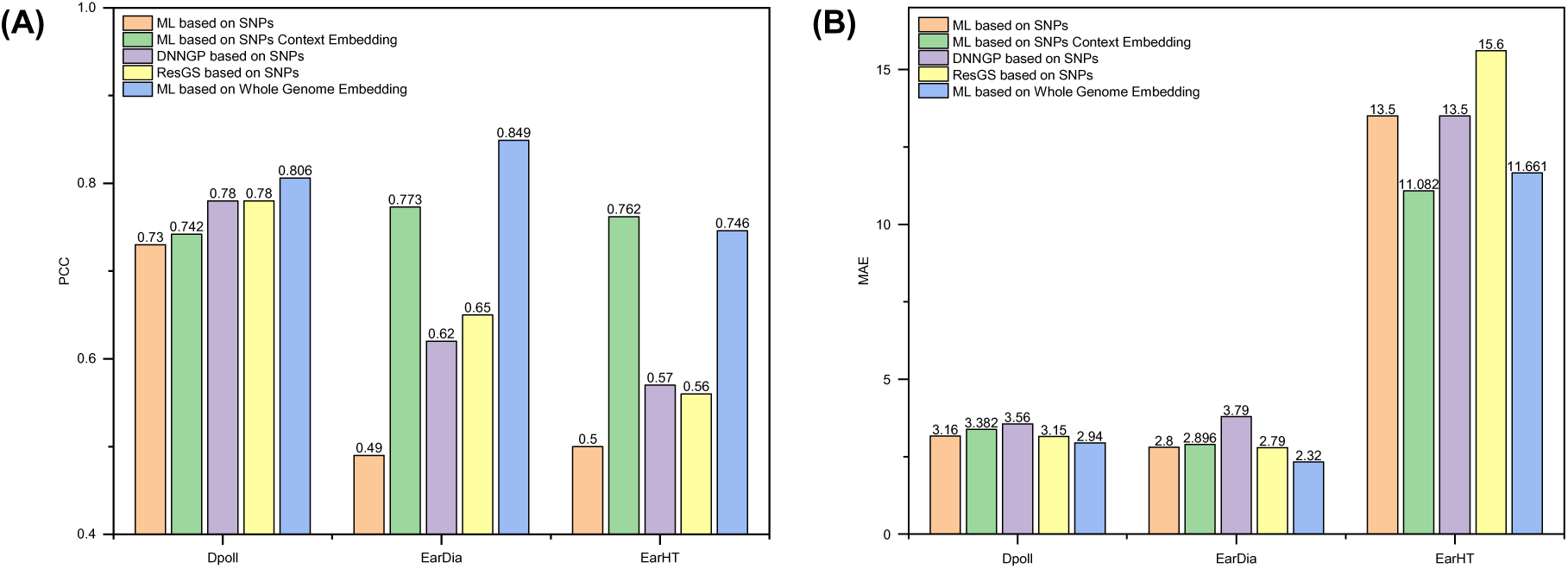
Performance comparison of traditional ML algorithms, which represent the type of algorithm that achieves optimal performance for each trait in Table 3 under different feature modes. DNNGP and ResGS are based on SNP markers, whereas ML algorithms are based on the SNP context and whole-genome feature embedding for traits in maize301. (A) PCC values for predicting Dpoll, EarDia, and EarHT traits. (B) MAEs for predicting the Dpoll, EarDia, and EarHT traits.

**Table 3.** Highest average prediction accuracy for various traits in maize301 under different feature modes.

| Traits for<br>maize301 | SNP* | SNP-context Feature<br>embedding |  |  | Whole-genome feature embedding |  |  |
| --- | --- | --- | --- | --- | --- | --- | --- |
|  | PCC | Algorithm<br>Type | Length of<br>Context | PCC | Algorithm<br>Type | Dimensionality<br>Reduction Method | PCC |
| Dpoll | 0.73 | SVR | 3000 bp | 0.742 | rrBLUP | PCA (d=16) | 0.806 |
| EarDia | 0.49 | rrBLUP | 3000 bp | 0.773 | GBR | NO_DR (d=768) | 0.849 |
| EarHT | 0.5 | RF | 2000 bp | 0.762 | rrBLUP | PCA (d=128) | 0.746 |

**Table 4.** Heritability of traits in rice413 based on genetic analysis.

| Trait for rice413 | Heritability |
| --- | --- |
| AC | 0.775324 |
| ASV | 0.412983 |
| PC | 0.417095 |
| PNPP | 0.485917 |
| SL | 0.982803 |
| SNPP | 0.505230 |

**Table 5.** Heritability of traits in rice395 based on genetic analysis.

| Traits for rice395 | Heritability |
| --- | --- |
| AC | 0.040703 |
| SL | 0.096478 |

**Table 6.** Heritability of traits in maize301 based on genetic analysis.

| Traits for maize301 | Heritability |
| --- | --- |
| Dpoll | 0.805676 |
| EarDia | 0.762109 |
| EarHT | 0.527156 |

**Table 7.** Average encoding time for three datasets based on two modes.

| Dataset | Embedding mode | Encoding Time Under Different Numbers of GPU(s) |  |  |  |
| --- | --- | --- | --- | --- | --- |
|  |  | 1 | 2 | 3 | 4 |
| Rice413 | SNP context (500) | 50.6 | 49.35 | 46.2 | 45.47 |
|  | SNP context (1000) | 67.46 | 65.62 | 64.56 | 62.96 |
|  | SNP context (1500) | 77.23 | 75.41 | 75.21 | 73.5 |
|  | SNP context (2000) | 79.47 | 76.9 | 75.84 | 75.05 |
|  | SNP context (2500) | 85.47 | 84.23 | 82.29 | 81.25 |
|  | SNP context (3000) | 92.64 | 92.22 | 91.73 | 89.18 |
|  | whole-genome embedding | 2006 | 1491.38 | 1108.98 | 969.87 |
| Rice395 | SNP context (500) | 5.95 | 4.09 | 3.73 | 3.2 |
|  | SNP context (1000) | 5.94 | 5.58 | 5.51 | 5.4 |
|  | SNP context (1500) | 6.02 | 5.81 | 5.45 | 5.08 |
|  | SNP context (2000) | 5.87 | 5.66 | 5.29 | 5.07 |
|  | SNP context (2500) | 6.47 | 6.44 | 5.76 | 5.08 |
|  | SNP context (3000) | 6.17 | 5.42 | 5.06 | 4.74 |
|  | whole-genome embedding | 1515.73 | 1310.42 | 1136.6 | 940.9 |
| Maize301 | SNP context (500) | 8.61 | 8.12 | 7.7 | 7.25 |
|  | SNP context (1000) | 10.61 | 10.07 | 9.57 | 9.08 |
|  | SNP context (1500) | 11.02 | 10.61 | 10.05 | 9.94 |
|  | SNP context (2000) | 11.48 | 11.32 | 11.14 | 10.56 |
|  | SNP context (2500) | 12.6 | 12.24 | 12.05 | 11.84 |
|  | SNP context (3000) | 12.94 | 12.53 | 12 | 11.51 |
|  | whole-genome embedding | 7214.67 | 6792.52 | 5879.26 | 4841.45 |

The significant performance enhancement for the EarDia trait using SNP-context feature embedding (3000 bp context length) and further improvement with whole-genome feature embedding suggest the presence of distal regulatory elements (>3000 bp from SNP sites) influencing trait expression. The high heritability of the Dpoll and EarDia traits in maize301, combined with the superior performance of whole-genome feature embedding, underscores its effectiveness in capturing genetic features associated with high-heritability traits. This aligns with observations in rice413, where whole-genome feature embedding excelled in predicting high-heritability traits such as SL and AC. Conversely, for medium-heritability traits such as EarHT in maize301 and ASV/PC in rice413, SNP-context feature embedding demonstrated comparable or superior performance to whole-genome embedding, highlighting its ability to uncover genetic interactions within localized genomic contexts.

### The fitting effect of typical traits under the two embedding modes

To compare the correlation and distribution of predicted versus observed values from whole-genome and SNP-context feature embedding, we selected one representative trait from each of three crops: EarDia (maize301), AC (rice395), and SL (rice413). We visualized the statistical distributions of the predicted and observed values via boxplots and normal distribution curves. Additionally, we perform regression analysis to derive the best-fit equations and visualize the linear regression results.

Figure 12 presents the regression performance for these three traits when both embedding modes are used. For the EarDia trait in maize301, whole-genome feature embedding produced predicted values with a median closer to the distribution of observed values, achieving the highest prediction accuracy. The regression fit exhibited narrow 95% confidence and prediction bands, with most data points aligning closely with the best-fit line. A high Pearson correlation coefficient (PCC ≈ 0.86) and goodness-of-fit (R²≈ 0.744) indicate robust model performance, likely due to the method’s ability to capture multidimensional genomic features such as structural variations and epigenetic modifications.

**Figure 12.**
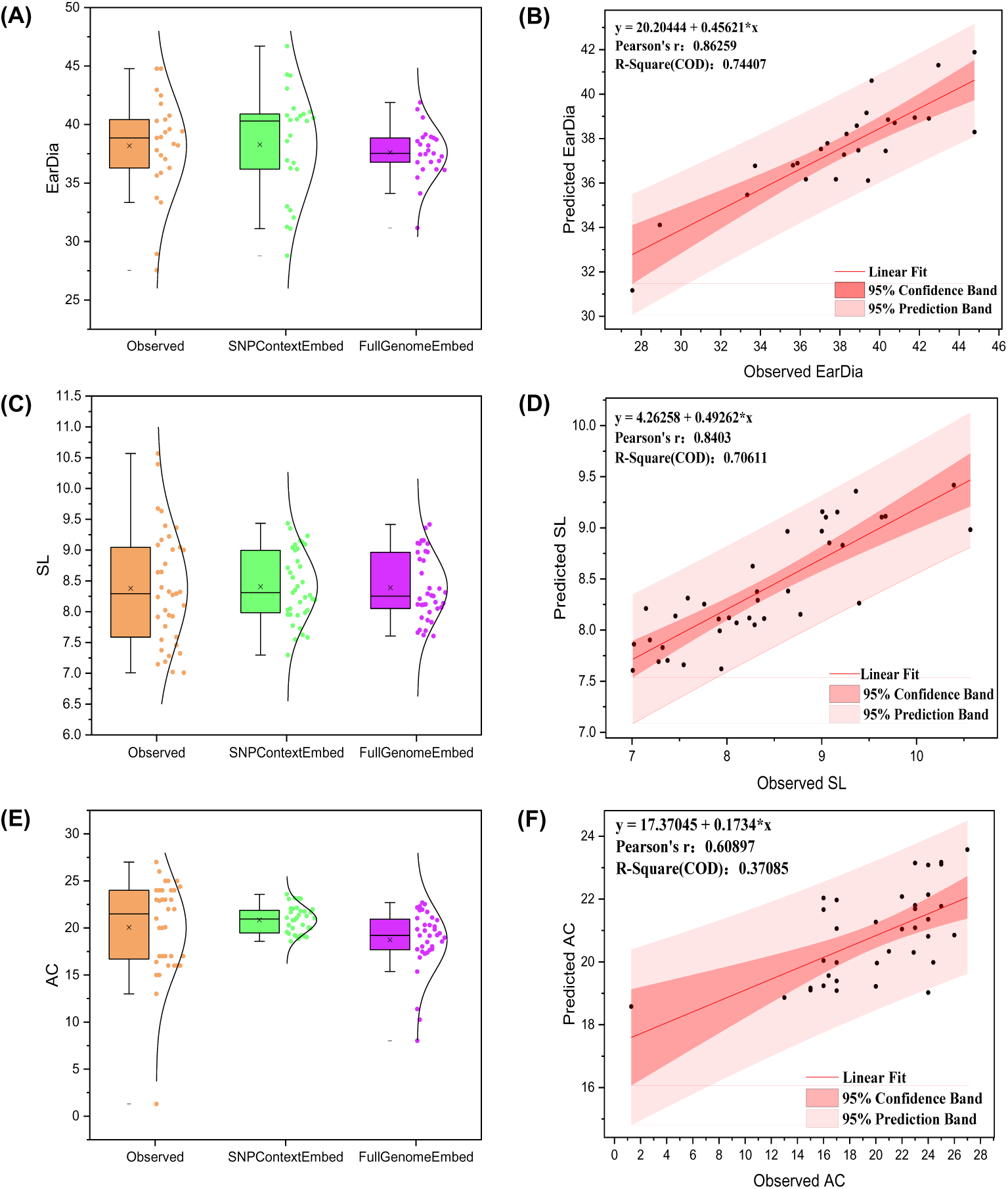
Fitting performance of typical traits under two embedding modes. (A, C, E) Boxplots and statistical distribution curves showing the predicted and observed values of the EarDia (maize301), SL (rice413), and AC (rice395) traits under SNP-context feature embedding and whole-genome feature embedding. (B, D, F) Fitting results between the predicted and true values for the EarDia, SL, and AC traits under optimal embedding conditions (as shown in Tables 1--3).

For the SL trait in rice413, whole-genome feature embedding also yielded predicted values with a median closer to the observed distribution. The fitting results showed narrow confidence and prediction bands, with data points concentrated around the best-fit line (PCC ≈ 0.84, R²≈ 0.71), reflecting stable and strong predictive capability.

In contrast, for the AC trait in rice395, SNP-context embedding was slightly more accurate than whole-genome embedding, but the overall prediction accuracy for both methods remained low. The wide 95% confidence and prediction bands, along with a low goodness-of-fit (R²≈ 0.37), indicate high prediction uncertainty. This suggests that traits with very low heritability, such as AC in this dataset, are highly susceptible to environmental influences, making it challenging for any model to capture their key genetic determinants.

### Comparative Analysis of the Encoding Times for the Two Modes

Whole-genome feature encoding is computationally intensive, particularly for large-scale genomic datasets. To address this challenge, we implemented a parallel processing strategy leveraging multi-GPU environments. We preprocessed complete crop genomes into 512-token sequence chunks to match the DNABERT-2 model’s input format, grouping them into batches of 100. Using data parallelism, these batches were distributed across available GPUs for accelerated encoding. Each GPU independently performs forward propagation on its allocated chunks to compute feature embeddings. Because the encoding of each chunk is independent, this approach minimizes the communication overhead and achieves highly efficient parallelization.

We calculated the average encoding time for whole-genome features and for SNP contexts of varying lengths (SNP context mode) in samples from the rice413, rice395, and maize301 datasets under different GPU configurations (as shown in Table 7). Across all three datasets, the average encoding time for whole-genome features was considerably longer than that for SNP contexts when the same number of GPUs was used. For the same crop type, as the number of GPUs increased, the average encoding time for whole-genome features progressively decreased. In the SNP context encoding mode, under the same GPU configuration, the overall average encoding time tended to increase as the SNP context length increased. Conversely, for a given SNP context length, increasing the number of GPUs resulted in some reduction in the average encoding time. It is evident that the time for whole-genome feature encoding progressively decreases as the number of GPUs increases. We believe that with a sufficient number of GPU resources or by leveraging high-performance computing (HPC) clusters with greater computational power, the time required for whole-genome feature encoding can be substantially reduced, potentially even to a level that is entirely acceptable for practical breeding applications.

## Discussion

In modern crop breeding, accelerating the development of elite varieties with superior agronomic traits and economic value is a central objective. As a revolutionary breeding strategy, genomic selection (GS) has significantly shortened breeding cycles and increased selection intensity when genome-wide markers are used to predict the genetic potential of individuals. Previous research has shown that many factors influence the accuracy of genomic prediction, such as the density and distribution of genetic markers, population size and structure, trait heritability, and gene‒environment interactions, among others [61, 62]. While SNP marker features, being the most common and abundant type of genetic variation, are utilized by most genomic prediction algorithms, they possess inherent limitations. For example, commercial SNP chips typically prioritize medium-to-high frequency SNPs, leading to the oversight of many rare variants with significant impacts on complex traits, and these marker sets do not encompass all types of genomic variation. Moreover, another notable limitation is that SNPs are often treated as independent markers, and the rich regulatory context of surrounding sequences and complex genetic interactions are easily overlooked. These problems can lead to an incomplete capture of the genetic architecture, particularly for traits influenced by nonadditive effects, rare variants with large effects that are poorly tagged by common SNPs.

This study utilized SNP contexts of varying lengths and whole-genome sequences as distinct inputs for the cross-species genomic foundational model DNABERT-2, constructing vector representations of SNP neighborhood genetic information and global genomic patterns. In addition, we explored the impact of different context lengths on four classical phenotype prediction algorithms. Our results demonstrate that ML algorithms operating under the SNP-context embedding mode achieve superior predictive performance compared with traditional SNP features at specific context lengths. Particularly for traits with low-to-moderate heritability (h^2^∈(0.2, 0.7], e.g., PNPP, SNPP, AC, ASV, and PC in rice413; EarHT in maize301), this advantage is most pronounced: nonlinear models (SVR and GBR) employing SNP-context embeddings exhibit significantly better performance than the linear rrBLUP model. Similarly, using whole-genome embedding as input can further improve the prediction accuracy for highly heritable traits (h^2^∈(0.7, 1.0], e.g., AC, SL in rice413; Dpoll, EarDia in maize301). The most compelling evidence comes from ML algorithms employing whole-genome features, which outperform even state-of-the-art deep learning models (such as DNNGP and ResGS) that rely on SNP marker features. This phenomenon may be attributed to the unique advantage of whole-genome feature encoding in characterizing complex, distributed genetic architectures—involving polygenic interactions, QTLs, and long-range genetic associations—which are typically undetectable by SNP-only or limited localized context, especially when genetic information serves as the predominant determinant of crop phenotypic traits.

While our study demonstrated the advantages of SNP-context feature embedding, it was limited to exploring context lengths within a 3000 bp range. We speculate that additional genetic features beyond this range may contribute to trait prediction, particularly for traits influenced by distal regulatory elements. Future research should investigate the impact of more extensive genomic contexts, potentially exceeding 3000 bp, to further increase the prediction accuracy. Moreover, a critical observation from our study is the diminished performance of both SNP-context and whole-genome embedding modes for traits with ultra-low heritability (h^2^∈(0, 0.2], e.g., AC, SL in rice395). In this scenario, DNNGP and ResGS with traditional SNP markers yielded better results. This aligns with established knowledge that traits with ultra-low heritability are largely influenced by environmental factors and complex gene‒environment (GxE) interactions, making their genetic signal inherently weak and difficult to capture consistently [27, 60]. The reduced genetic stability across generations for such traits means that even sophisticated genomic models struggle to disentangle true genetic contributions from environmental noise. To address this problem, it is indeed feasible and highly promising to integrate multi-omics data to improve the prediction of ultra-low heritability traits. Environmental influences often manifest through changes at various molecular levels. Incorporating data from transcriptomics (gene expression), epigenomics (e.g., DNA methylation), proteomics, or metabolomics could provide a more dynamic and comprehensive picture of an organism’s state. These data layers are often closer to the phenotype and can capture the molecular consequences of environmental perturbations or GxE interactions. For example, gene expression levels can directly reflect how an organism responds to environmental stress, and integrating this information could help partition out environmental effects or model GxE explicitly, thereby enhancing the accuracy of predictions, especially for traits where the direct genomic signal is weak. Future research should explore multi-modal learning architectures that can effectively fuse genomic embeddings with these other omic layers.

Our method provides new insights into the precise prediction of complex traits, demonstrating both theoretical significance and practical value in accelerating crop improvement. However, current GPU computational limitations impose substantial time requirements for encoding crop whole-genome features in our experiments. For widespread implementation in large-scale breeding programs, future advancements must address three key challenges: (1) adopting more advanced GPUs with increased core counts and VRAM capacity through high-performance computing clusters or cloud-based elastic computing resources; (2) exploring model compression techniques including knowledge distillation (training smaller models to replicate DNABERT-2’s embeddings), pruning, and quantization to reduce model size and computational demands while maintaining predictive accuracy; and (3) developing more efficient data and model parallelism strategies with optimized distributed training frameworks to maximize multi-GPU/multi-node utilization and reduce encoding time.

## Data availability

The authors affirm that all the data necessary for confirming the conclusions of the article are present within the article/supplementary material. Further inquiries can be directed to the corresponding author.

## Supplementary information

Supplementary material is available at *Plant Methods online*.

## Acknowledgments

This work was supported by the Open Fund of the Key Laboratory of Agricultural Big Data, Ministry of Agriculture and Rural Affairs (No. DSJSYS-2024-02), the Central Public-interest Scientific Institution Basal Research Fund (No. JBYW-AII-2025-21)

## Author contributions

HL finished the experiment and wrote the manuscript. TW, CW and WB assisted in collecting phenotypic data and genomic data and restoring whole-genome sequences. JL and LC played crucial roles in data collection, participating in the preprocessing of phenotypic and genomic sequences. YC and ZC provided expert guidance in data analysis and offered invaluable feedback on the manuscript. MWs jointly supervised the entire research process, ensuring ethical compliance and effectively facilitating communication among authors. Finally, JW and JH were responsible for proofreading the manuscript. All authors reviewed the manuscript and approved the final manuscript.

## Conflict of interest

The author(s) declare no conflicts of interest.

## Notes

### Competing Interest Statement

The authors have declared no competing interest.

### Summary of Updates

Section on Method implementation added Figure 1 D Section on Results added Comparative Analysis of the Encoding Times for the Two Modes Section on Discussion updated

